# Application of CMSY method to estimate biological reference points of Piracatinga (*Calophysus macropterus* Lichtenstein, 1819) from the upper Solimões River in the Amazon Basin

**DOI:** 10.1101/2025.10.20.683447

**Authors:** Alfredo Pérez Lozano, Donald Charles Taphorn

**Affiliations:** Universidade Federal do Amazonas-UFAM, Programa de Pós-graduação em Ciência e Tecnologia para Recursos amazônicos, Itacoatiara, AM, Brasil; Universidad Nacional Experimental de los Llanos Occidentales Ezequiel Zamora (UNELLEZ), BioCentro, Guanare, Portuguesa, Venezuela

**Keywords:** Fisheries resources, Stock management, Overfishing, Data Limited Fisheries, Brazil

## Abstract

The Amazon basin contains a wide variety of commercially valuable fish species, with fishing being the main source of income for artisanal fishermen and riverside communities. After the decline in large catfish production in the late 1990s, Piracatinga emerged as an alternative to the local economy and was intensively exploited between 1996 and 2012. However, its capture became uncontrolled and was finally banned in 2015. Despite this, an assessment has never been conducted. The objective of this study was to determine Biological Reference Points (BRP) to estimate the level of exploitation. For this purpose, the (CMSY) assessment method was applied, using fishing statistics. The results showed that Piracatinga is in a sustainable situation. The biomass indicators for optimal sustainable yield (B_MSY_ = 2.25 t) and fishing mortality (F) values at the maximum sustainable yield point (F_MSY_ = 0.55). Therefore, the population status of the Piracatinga stock is in “good condition” (F_2012_/F_MSY_ (<1) and B_2012_/B_MSY_ (>1). These results indicate that despite the good condition of the resource, we recommend precautionary measures of a technical and legal nature before allowing its exploitation again.

## 1. INTRODUCTION

The Amazon Basin is one of the largest hydrographic regions in the world, characterized by a wide diversity of habitats that sustain rich fauna and flora, making it one of the most biodiverse regions on the planet. This basin spans six countries: Brazil, Peru, Bolivia, Colombia, Ecuador, and Venezuela (Fabré & Alonso 1998, Agudelo et al. 2000). Among the region’s fishery resources, species of the family Pimelodidae have significant economic importance, including the Gilded Catfish (*Brachyplatystoma rousseauxii*), the Tiger Shovelnose Catfishes (*Pseudoplatystoma punctifer* and *Pseudoplatystoma tigrinum*), the Jau Catfish (*Zungaro zungaro*), the Pirarara (*Phractocephalus hemioliopterus*), and the Piraíba (*Brachyplatystoma filamentosum*), However, many aspects of their life cycles, population dynamics, and exploitation status remain poorly understood or largely unknown (Agudelo et al. 2000, Petrere et al. 2004, Deza-Taboada et al. 2006, Agudelo et al. 2013, García et al. 2013, Freitas & Montag 2019, de Oliveira-Diaz et al. 2024).

During the 1990s, intensive commercial over-fishing of large catfish species in the Upper Solimões led to a significant shift in catch composition. The drastic decline of these species resulted in their replacement by smaller, lower-value species, which had previously been underrepresented in official fishery statistics (Petrere et al. 2004). Within this context, Piracatinga fishing began gaining importance in 1997 in the Amazon River (Colombian Amazon), driven by the collapse of large pimelodid catfish stocks and the decline in catches of *Pimelodus grosskoppii*, a species highly valued in Colombia (Franco et al. 2016). By 2001, Piracatinga (*Calophysus macropterus* Lichtenstein, 1819) had already acquired considerable commercial relevance, meeting an increasing market demand. In the tri-border region of Peru, Colombia, and Brazil alone, the harvest reached 1,300 t (Table I).

**Table I.**
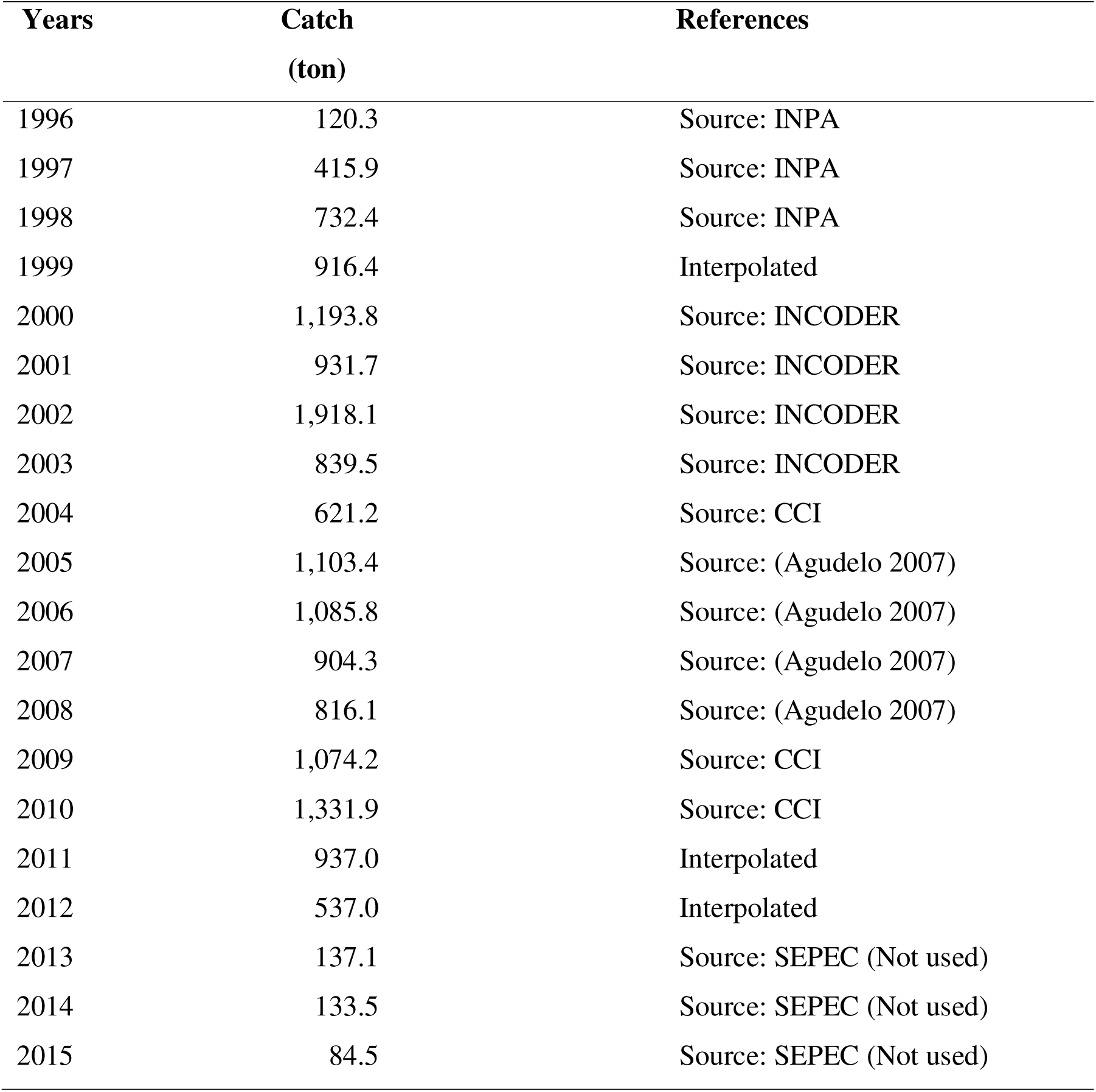
Annual production (in tons) of Piracatinga in Leticia, Colombia, adapted from records of Colombian Fishing Statistics Yearbooks (INPA, CCI, INCODER, SEPEC) and Agudelo (2007).

| Years | Catch<br>(ton) | References |
| --- | --- | --- |
| 1996 | 120.3 | Source: INPA |
| 1997 | 415.9 | Source: INPA |
| 1998 | 732.4 | Source: INPA |
| 1999 | 916.4 | Interpolated |
| 2000 | 1,193.8 | Source: INCODER |
| 2001 | 931.7 | Source: INCODER |
| 2002 | 1,918.1 | Source: INCODER |
| 2003 | 839.5 | Source: INCODER |
| 2004 | 621.2 | Source: CCI |
| 2005 | 1,103.4 | Source: (Agudelo 2007) |
| 2006 | 1,085.8 | Source: (Agudelo 2007) |
| 2007 | 904.3 | Source: (Agudelo 2007) |
| 2008 | 816.1 | Source: (Agudelo 2007) |
| 2009 | 1,074.2 | Source: CCI |
| 2010 | 1,331.9 | Source: CCI |
| 2011 | 937.0 | Interpolated |
| 2012 | 537.0 | Interpolated |
| 2013 | 137.1 | Source: SEPEC (Not used) |
| 2014 | 133.5 | Source: SEPEC (Not used) |
| 2015 | 84.5 | Source: SEPEC (Not used) |

In Colombia, between 1996 and 2004, the reported Piracatinga catch volume increased by 92.81% compared to initial values (Gómez et al. 2008). Subsequently, its production expanded significantly in the mid-Solimões region (Franco et al. 2016), and fishing activities were further extended into the western Brazilian Amazon in 2000 (Brum et al. 2015, Iriarte & Marmontel 2013). Later, Piracatinga fishing reached Bolivia in the Mamoré River basin, where fishing techniques previously developed in Brazil and Colombia were adopted (Van Damme et al. 2023).

In Peru, Piracatinga landings between 2006 and 2016 ranged from 7,393 to 4,247 t., showing a slight downward trend (Del Águila-Chávez et al. 2023). Reports from 2007 indicate that Brazil produced 1,600 t. of Piracatinga (Barthem and Goulding, 2007, Brum et al. 2015). By 2011, production in the Amazonas state, Brazil, had reached approximately 4,400 t. (Amazonas State Production Secretariat, unpublished data, *apud*. Brum et al. 2015).

However, due to negative media coverage regarding the use of river dolphins (*Inia geoffrensis*) and caimans (*Caiman crocodilus*, *Melanosuchus niger*) as bait for Piracatinga fishing, the Brazilian government enacted a five-year moratorium in 2015 (Ministério do Meio Ambiente, 2014). The Brazilian government implemented this measure, promising that fishery resource assessments would be conducted during this period to support the development of a management plan. Nonetheless, the capture of this species remains prohibited to this day.

The piracatinga, like other small commercial marine species such as pink conger eel (*Genypterus brasiliensis*), has excellent-quality meat and is of great gastronomic interest (Tomas et al. 2019). Unfortunately, as with the pink conger eel, there is no available data on fishing effort for the piracatinga, which would allow estimates of the resource’s status for Brazilian fisheries.

Furthermore, despite the unregulated exploitation of this fishery, few studies have investigated its bioecology and fishery dynamics (Pérez and Fabré 2002, 2009, Garcia et al. 2017, Bonilla et al. 2022). To date, only Bonilla et al. (2022) have estimated the exploitation rate of Piracatinga using the expression E = F/Z (Gulland 1969). However, no studies using stock assessment methods have determined the population status of this species, either unilaterally or through coordinated efforts among the countries sharing this fishery.

Tropical commercial fisheries are notoriously difficult to manage due to the scarcity of catch and effort data. Small-scale fisheries are often marginalized due to the lack of reliable information, making it challenging to implement effective management strategies and allowing overfishing to go unnoticed until its impacts on stocks and fishing communities become evident (Pauly 1997, Sparre & Venema 1997). Therefore, the development of alternative approaches that do not rely exclusively on traditional quantitative data is necessary (Johannes 1998). In recent years, advancements in stock assessment methods for data-limited fisheries have helped bridge this gap, enabling the identification of overfishing signals and population declines even when available data are scarce (Pereira & Hansen 2003, Prince et al. 2015, Pons et al. 2020).

The FAO (1995) recommends the use of Biological Reference Points (BRPs) as a tool to assess the sustainability of fishery resources. Scientists have developed mathematical methods based on different approaches, such as catch-based (Berkson et al. 2011, Newman et al. 2015, Carruthers et al. 2014), abundance-based (Froese et al. 2022), and length-based distribution models (Froese 2004, Cope & Punt 2009, Hordyk et al. 2015), to support stock assessments. Among the most widely used methods for stock assessment in data-limited fisheries are CMSY (Catch-MSY), developed by Froese et al. (2017), which utilizes retrospective catch data and species resilience. This method is fundamental for evaluating the sustainability of Piracatinga fisheries.

Given this context, the present study aims to determine the BRPs for Piracatinga in the Upper and Middle Solimões using historical catch data from 1996 to 2012 and applying data-limited stock assessment methods. By doing so, this research seeks to establish a baseline for the management of this resource and contribute to a differentiated approach to stock assessment in the Amazon region.

## 2. MATERIAL AND METHODS

### 2.1. Study Area

The Upper Solimões region is located approximately between latitudes 0° and 4° S and longitudes 63° and 73° W, within the tri-border area shared by Peru, Colombia, and Brazil. The Piracatinga fishing grounds are located in the Peruvian and Colombian portions of the Upper Solimões (Chimbote, Puerto Alegría, Puerto Nariño, Santa Rosa, Leticia) and in the Brazilian portion of the Middle Solimões (Benjamin Constant, Atalaia do Norte, Jutaí, Fonte Boa, Tefé, and Coari) (Figure 1). Throughout the period from 1996 to 2012, Piracatinga production was concentrated in Leticia (Colombia) and partially in Tabatinga (Brazil).

**Figure 1.**
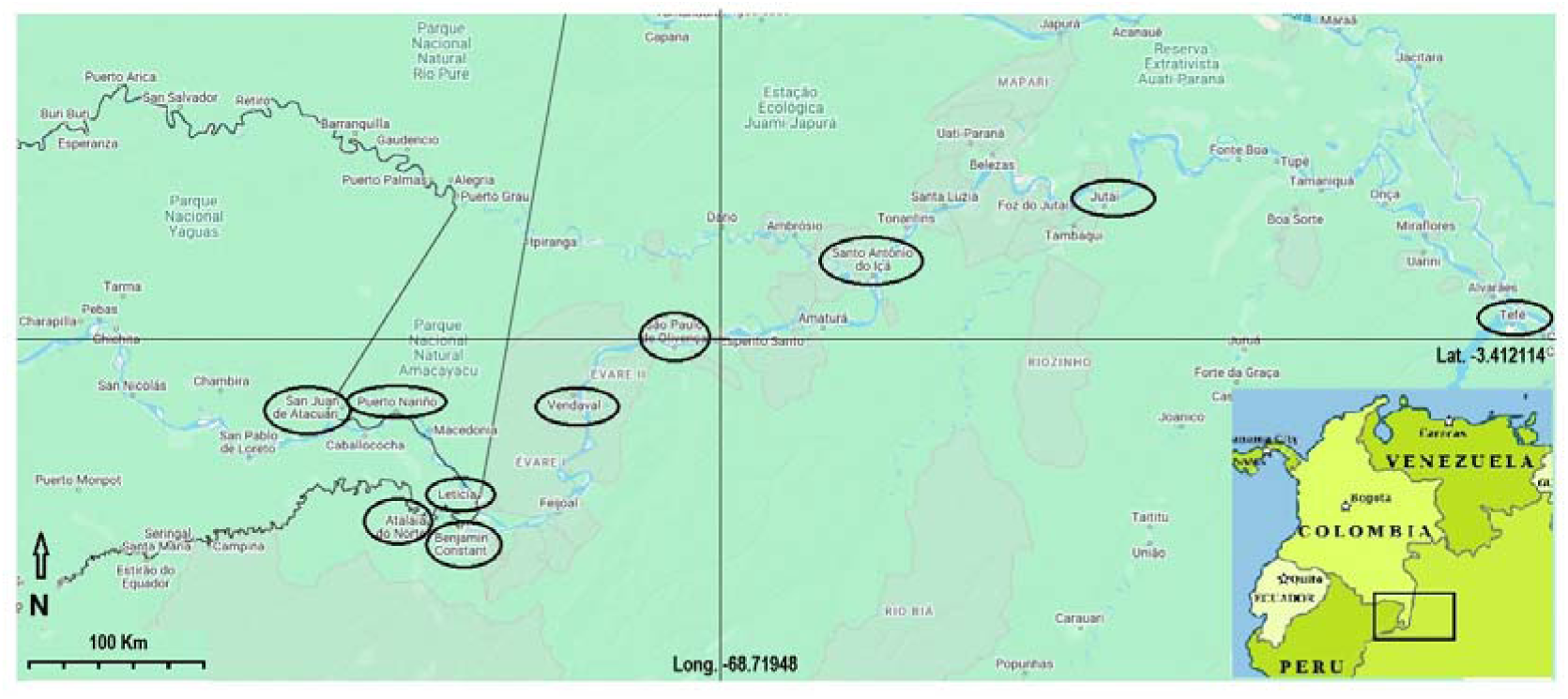
Map of the main Piracatinga fishing areas in the Upper and Middle Solimões in Brazil, Colombia and Peru.

### 2.2. Available data

For the analysis of Piracatinga catch data, a 17-year historical series (1996–2012) was used. These data were collected from various Colombian Fishing Statistics Yearbooks: Instituto Nacional de Pesca y Acuicultura (INPA), Instituto Colombiano de Desarrollo Rural Pesquero (INCODER), Corporación Colombiana Internacional (CCI), Servicio Estadistico Pesquero Colombiano (SEPEC). All catch data, originally recorded in kilograms, were converted to metric tons (t).

### 2.3. Catch-MSY method

The CMSY, is a method developed by Martell & Froese (2013) and improved by Froese et al., (2017) to estimate Maximum Sustainable Yield (MSY). This approach is based on Schaefer’s (1954) surplus production model and utilizes species-specific catch and resilience data. It considers the parameters r (intrinsic growth rate) and k (carrying capacity), where different combinations of these parameters generate various biomass time series. So, CMSY is a very simple and, therefore, practical tool (Froese et al. 2019).

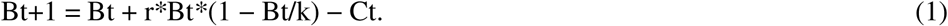

Where: Bt= biomass at time step t of the time series; r= intrinsic growth rate; k= environmental carrying capacity (assumed equal to the unexploited biomass B□); Ct= known catch at time stept.

The CMSY method models population dynamics based on production and incorporates information on species resilience, allowing for parameterization of the intrinsic population growth rate (r) and the carrying capacity (k) of the unexploited population. The estimation of viable r-k pairs is performed using an iterative Markov Chain Monte Carlo (MCMC) approach. Additionally, the model does not assume negative values and is consistent with the prior definition of relative biomass values at the beginning and end of the time series (Froese et al. 2019).

After identifying an optimal pair of r and k, a biomass time series (B) and fishing mortality (F) can be calculated, along with several key indicators (Froese et al. 2016), as follows:

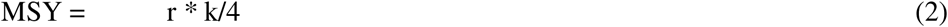

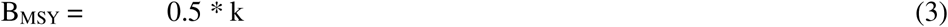

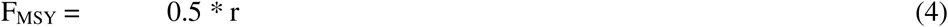

Thus, the biological reference points of the stocks can be defined based on the B/B_MSY_ and F/F_MSY_ ratios from the final year of a time series (Table II).

**Table II.**
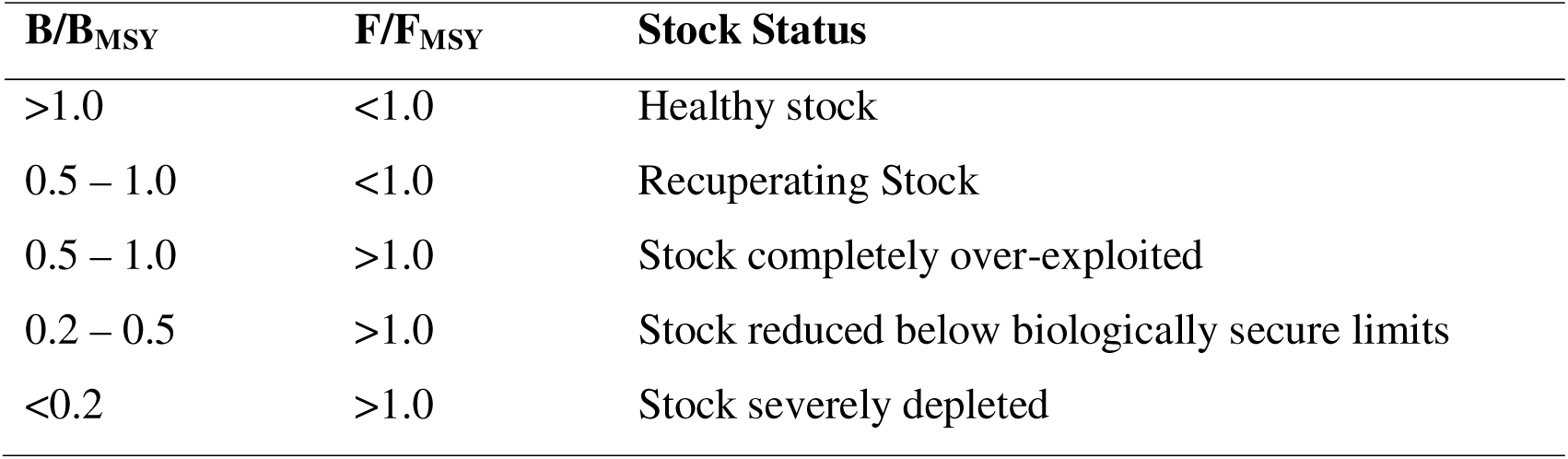
Definition of fish stock status for fisheries management based on B/B_MSY_ and F/F_MSY_ in the final year of a time series (Froese et al. 2016).

| $B/B_{MSY}$ | $F/F_{MSY}$ | Stock Status |
| --- | --- | --- |
| >1.0 | <1.0 | Healthy stock |
| 0.5 – 1.0 | <1.0 | Recuperating Stock |
| 0.5 – 1.0 | >1.0 | Stock completely over-exploited |
| 0.2 – 0.5 | >1.0 | Stock reduced below biologically secure limits |
| <0.2 | >1.0 | Stock severely depleted |

### 2.4. Input Parameters and Data

The input parameters used in the model include species resilience and relative biomass (B/k) intervals corresponding to depletion levels at the initial, intermediate, and final time series points. Resilience is a preliminary estimate of a species’ ability to absorb intrinsic and extrinsic population impacts, expressed through intrinsic growth rates. The suggested values are categorized as “high,” “medium,” “low,” and “very low.” Table III presents the r intervals automatically assigned by the CMSY model based on resilience categories, sourced from www.FishBase.org.

**Table III.** Resilience categories and their corresponding “r” values (Froese et al. 2019).

| Resilience | r – Low | r – High |
| --- | --- | --- |
| High | 0.600 | 1.500 |
| Mean | 0.200 | 0.800 |
| Low | 0.050 | 0.500 |
| Very Low | 0.015 | 0.100 |

Moreover, it is common for some species to exhibit different resilience values across distinct populations. Therefore, it is possible to take advantage of the relationship between r and other parameters, such as:

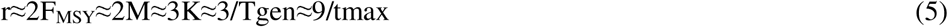

Where: r = intrinsic population growth rate; F_MSY_= fishing mortality rate at MSY (0.5*r); M= natural mortality rate; K= von Bertalanffy growth function (VBGF) parameter; Tgen= generation time of the population; Tmax= maximum age of individuals in the population (Froese et al. 2017).

To define the relative biomass (B/k) intervals used in the CMSY method, we applied the reference values provided by Froese et al. (2019) (Table IV).

**Table IV.**
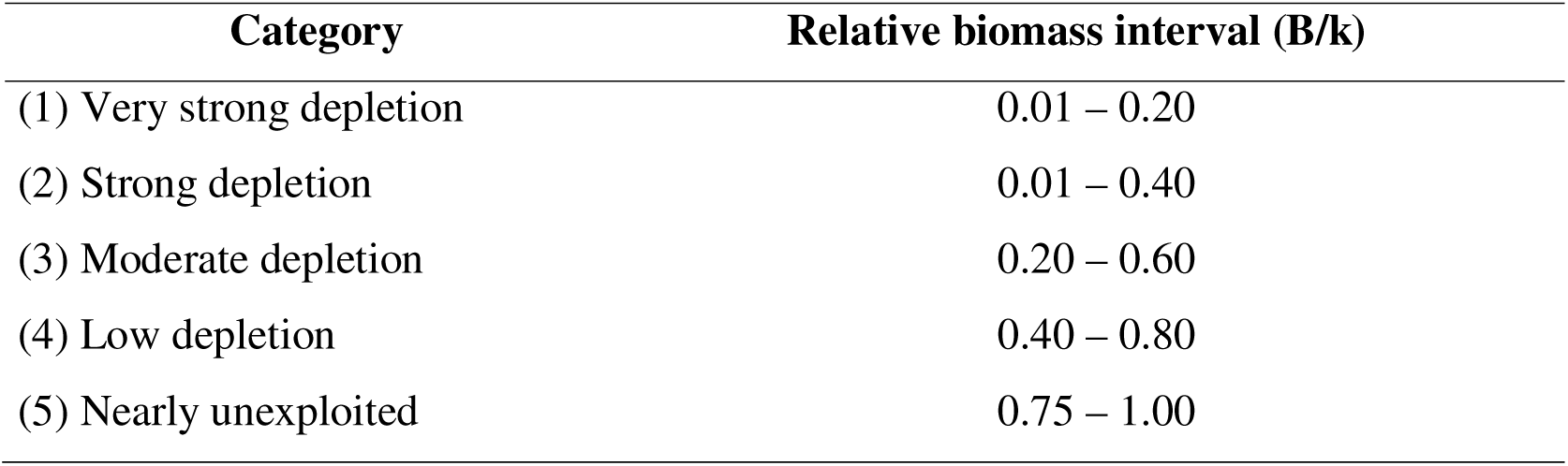
Depletion levels based on relative biomass (B/k) intervals (Froese et al. 2019).

Based on the framework proposed by Drescher et al. (2013) and Martin et al. (2012) for incorporating missing formal knowledge, expert elicitation was used to define input values for the model. The selection of value pairs relied on informed judgment by the authors, who possess extensive expertise in Amazonian fisheries. The input parameters adopted for the CMSY model, applied to assess the population status of Piracatinga, are presented in Table V.

**Table V.** CMSY model input parameters for Piracatinga in the Upper Solimões River.

| **Parameter** | **(r–Low, r–High) Input Values** |
| --- | --- |
| Initial relative biomass (B/k) | 0.300 - 0.800 (expert judgment) |
| Intermediate relative biomass (B/k) 2003 | 0.349 - 0.800 (expert judgment) |
| Final relative biomass (B/k) | 0.402 - 0.794 (expert judgment) |
| Initial variation for r | 0.569 - 1.500 (expert judgment) |
| Initial variation for k | 2.450 - 12.30 |
| Time period (years) | 1996 – 2012 |

All statistical analyses were conducted using the R programming language (R Development Core Team 2021). The CMSY method was implemented using the script CMSY_2019_9f.R (https://github.com/SISTA16/cmsy), Described in Froese et al., (2017), using the iMarine platform (https://i-marine.d4science.org/group/sdg-indicator14.4.1/stock-monitoring-tools-v0.6), an initiative launched in 2015 to establish and operate an e-infrastructure supporting the ecosystem-based approach to fisheries management and the conservation of living marine resources, aligned with the FAO’s Blue Growth Initiative.

## 3. RESULTS

The results in Table VI indicate that the Piracatinga stock was in a state that can be considered “healthy.” This conclusion is supported by the stock biomass indicators for 2012 (B_2012_ = 3.10 t) and the biomass at maximum sustainable yield (B_MSY_ = 2.25 t). Additionally, the fishing mortality rate (F_2012_ = 0.30) and the fishing mortality at MSY (F_MSY_ = 0.54). These results suggest that the Piracatinga stock was in an “optimal state” (F_2012_/F_MSY_ = 0.55), remained below one (<1.00). Moreover the (B_2012_/B_MSY_=1.37), was above the reference threshold (>1.00), these findings indicate that the stock was indeed in a healthy condition.

**Table VI.**
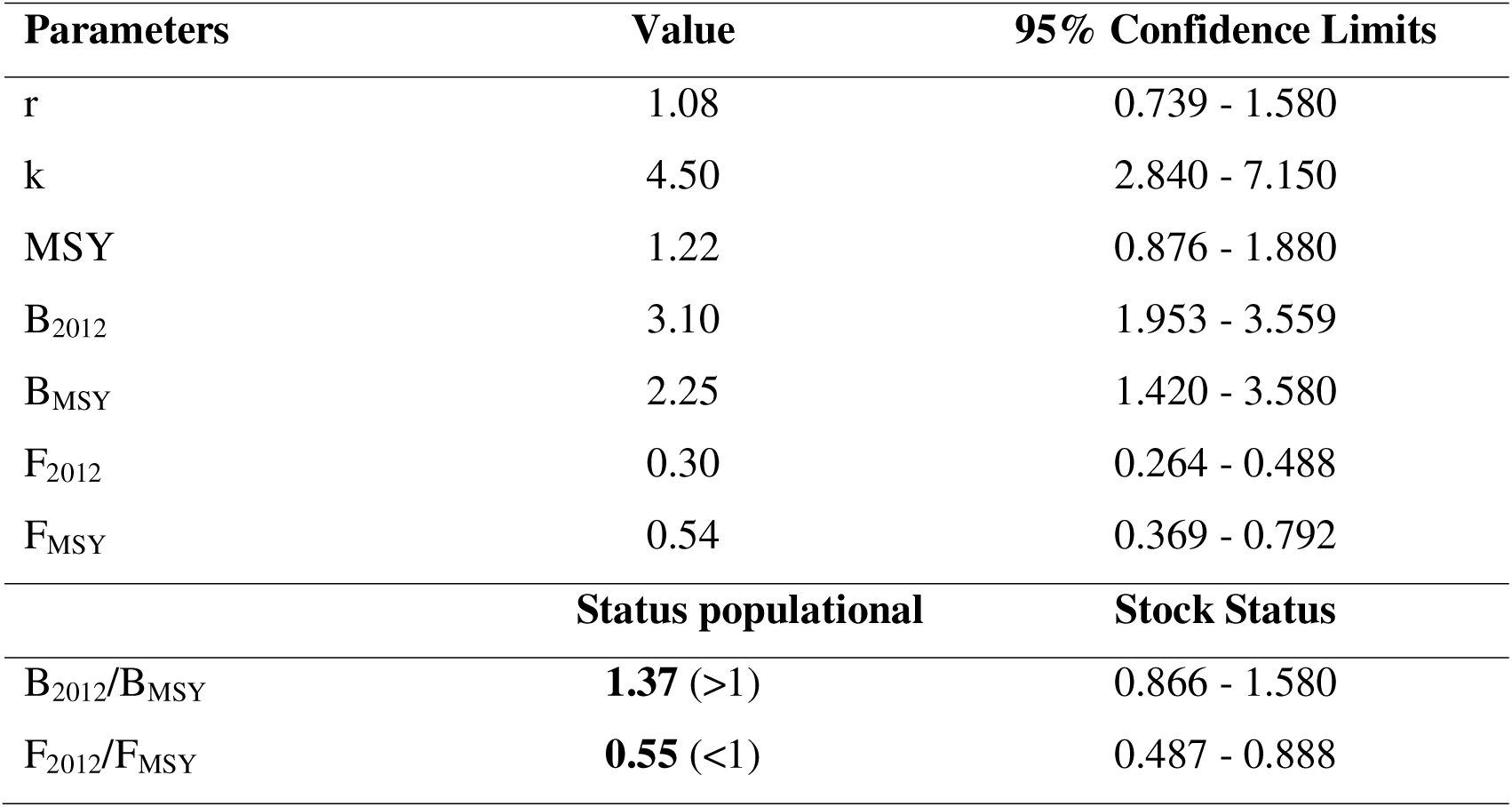
The results of the CMSY analysis are based on viable r–k pairs and results for management based on the CMSY analysis. And the stock status of Piracatinga determined from the combinations of the pairs B/B_MSY_ and F/F_MSY_

The fishing mortality results demonstrated that at the end of the historical data series analysed in 2012 (F_2012_ = 0.30), the value was significantly below the MSY threshold (F_MSY_ = 0.54). Regarding F/F_MSY_, a sharp upward trend was observed, crossing the sustainable threshold only in 2003 and reaching its peak that same year (Figure 2). However, the exploitation rate (F/F_MSY_) exceeded 1 during 2002–2003 (peak ∼1.5), then decreased to 0.55 in 2012; therefore, the status at the end of the series is <1, although there was episodic growth overfishing in the early 2000s.

**Figure 2.**
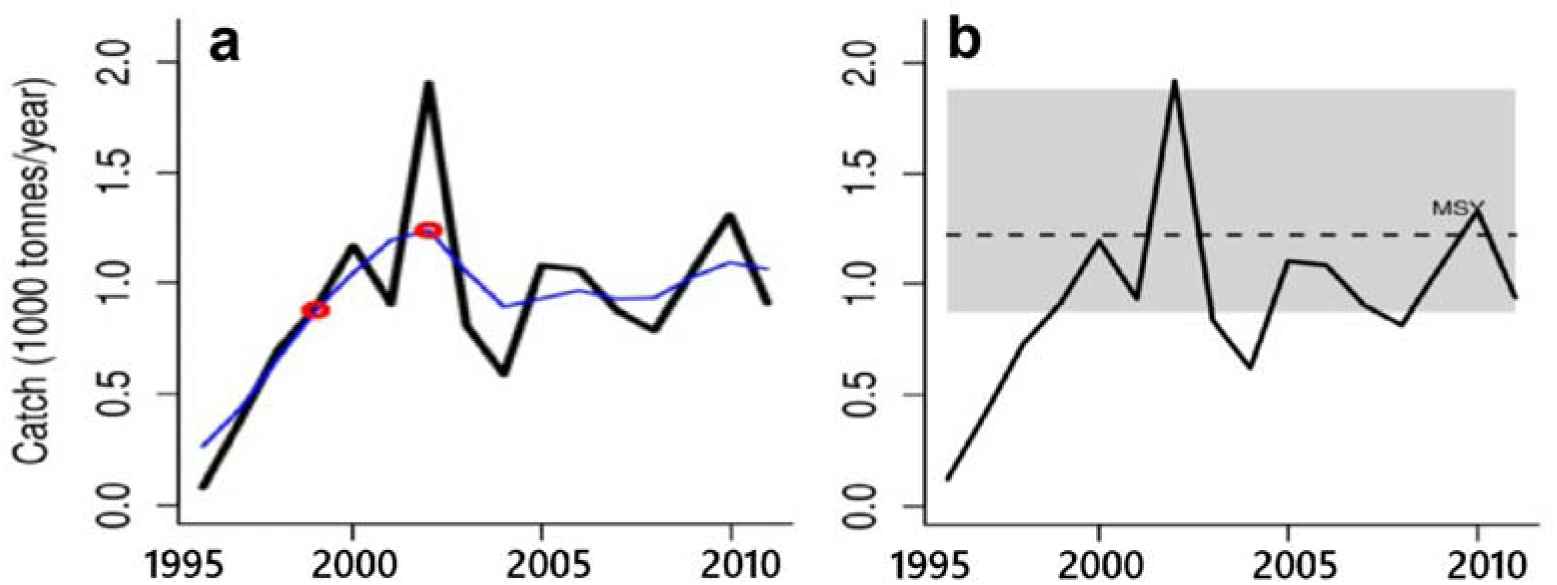
a) Time series of Piracatinga catches from 1996 to 2012, in the upper Solimões in black, with the thin blue curve representing smoothed catch and the red circles the smoothed minimum and maximum values. b) Temporal variation of catches over time compared to the MSY estimated in the CMSY, accompanied by a 95% confidence interval (shaded in grey).

The biomass pattern, influenced by fishing mortality and recruitment, peaked in 2002 and subsequently declined continuously until 2011, with further reductions observed in 2012 and finally in 2015, when Piracatinga fishing was halted. The catch and stock size graph indicates a transition phase in stock status, wherein the resource remained within a safe zone for several years and approached depletion only toward the end of the analysed period, without surpassing the F_MSY_ limit. The B/B_MSY_ trajectory in Figure 3 illustrates a progressive decline, with a notable decrease beginning in 2003 and intensifying thereafter, reaching a minimum value in 2012 but remaining above the target reference point (B/B_MSY_ = 1.00).

**Figure 3.**
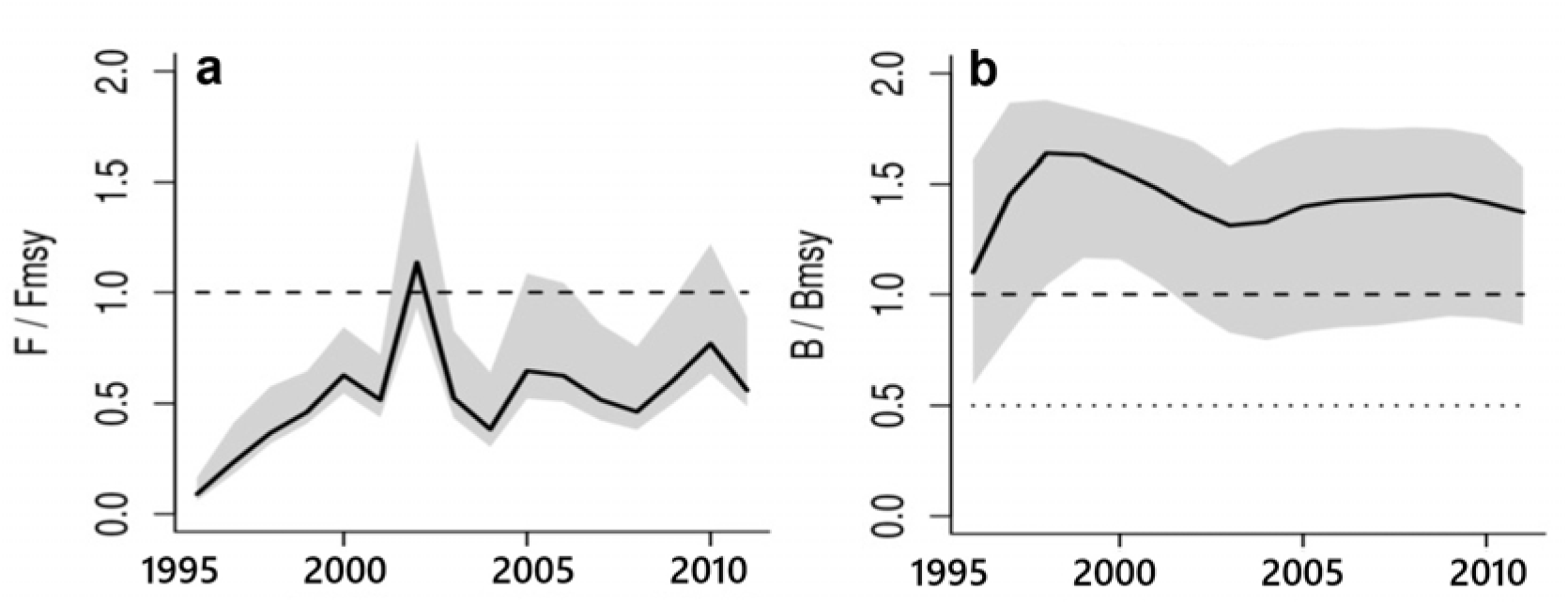
a) Temporal variation of relative fishing mortality (F/F_MSY_), with the trend of exploitation (F/F_MSY_), accompanied by a 95% confidence interval (shaded in grey). b) Temporal variation of the trend of relative biomass (B/B_MSY_), with the F_MSY_ value corrected for recruitment reduced below 0.5 B_MSY_, accompanied by a 95% confidence interval (shaded in grey).

The Kobe plot, which describes the simultaneous trajectory of B/B_MSY_ and F/F_MSY_ (Figure 4), demonstrates a progressive temporal trend of the resource, remaining within the green zone from 1996 to 2012. During this period, the Piracatinga stock never approached the red zone, passing briefly through the orange area, likely around 2003, when its maximum catch was recorded. The data indicate that despite intense fishing pressure, the stock never entered the red zone nor approached it toward the end of the dataset, suggesting that fishing never posed a risk of overexploitation.

**Figure 4.**
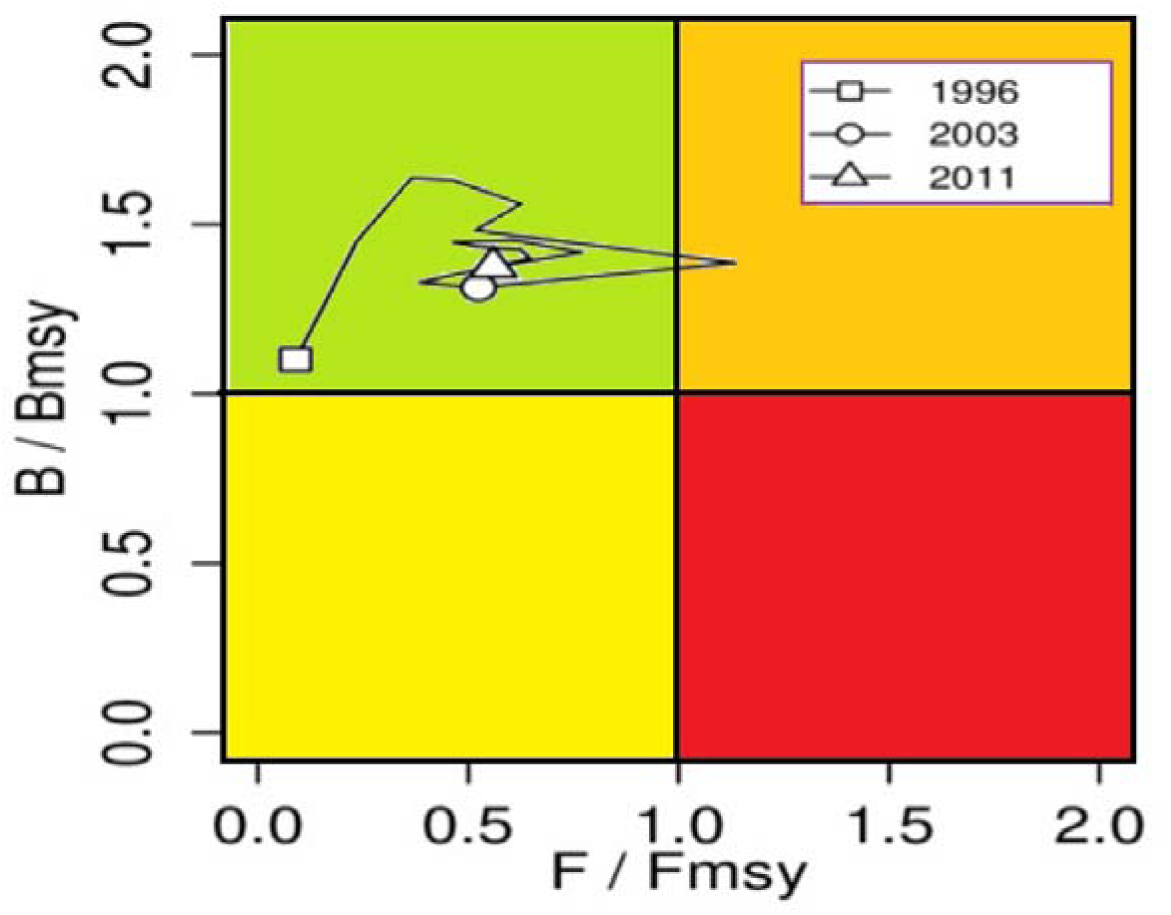
Kobe plot depicting simultaneous growth in exploitation (F/F_MSY_) and relative biomass (B/B_MSY_). The green area delineates the fishing safety zone, where MSY is achieved through sustainable fishing pressure and robust biomass; the orange zone signals imminent risk of overfishing as a result of increased fishing pressure on the stock biomass; the red area represents depleted stock biomass unable to generate MSY due to persistent overexploitation; and the yellow zone denotes the recovery stage characterized by decreasing fishing pressure.

The status of the Piracatinga stock, as evaluated by the CMSY method, is classified as good. The F/F_MSY_ indicator remained below 1 throughout the analysis period and has remained stable, while the B/B_MSY_ indicator has consistently been above 1 since 1996. The MSY value was estimated at approximately 1,220 t (95% C.I. = 876 – 1,880 t), with MSY being exceeded only in 2002 (1,918 t). Thus, based on the CMSY method, the Piracatinga stock in the Solimões River can be considered in a satisfactory condition.

## 4. DISCUSSION

The results obtained via the CMSY method indicate that the Piracatinga stock during the analysed period (1996–2012) was in a healthy state, with biomass values above recommended biological reference points (>1.00). The relationship between F_2012_/F_MSY_ and B_2012_/B_MSY_ further supports this conclusion, suggesting that the population was not overexploited during the assessed period. Additionally, a graphical analysis of the Kobe plot demonstrated that, despite relatively intense fishing pressure, Piracatinga did not reach critical levels of overexploitation.

However, some studies suggest that catch increases may reflect either heightened fishing efforts or structural changes in the exploited population (Bosch et al. 2021), underscoring the need for continuous monitoring to mitigate potential impacts on fishery sustainability.

Our findings contrast with those of Bonilla et al. (2011), Nuñez-Avellaneda et al. (2007), and Bonilla et al. (2022), who reported slower growth rates and higher fishing pressure on Piracatinga populations in the Putumayo River (a Solimões tributary)—a phenomenon termed “stock tropicalization” (Stergiou 2002). However, tropicalization typically applies to long-lived species subjected to selective fishing pressure, leading to earlier maturation and reduced average body size. Our results suggest that size differences observed by Bonilla et al. (2022) between the Putumayo (larger fish) and the Ucayali-Marañón confluence (smaller fish) may relate more to proximity to spawning grounds than solely to fishing intensity.

The reliability of data-limited methods like CMSY has been widely debated (Froese 2004, Froese et al. 2016, 2017). While useful for data-poor fisheries, their limitations are well-documented. For instance, CMSY tends to overestimate fishing mortality and underestimate biomass (Sharma et al. 2021). Reliance on official catch statistics may also introduce bias, as these often exclude subsistence fishing, IUU (Illegal, Unreported, and Unregulated) activities, and temporal reporting gaps (Gushchin & Shavrina 2018, Andrašunas et al. 2022).

A further challenge lies in CMSY’s heuristic models. In well-managed stocks, the method may erroneously interpret declining catches as increased fishing pressure, distorting data interpretation (Pons et al. 2020). This highlights the need for expert validation and multi-indicator approaches to minimize bias (Andrašunas et al. 2022).

While empirical, data-limited methods provide preliminary insights into stock status, they rarely offer definitive sustainability benchmarks. Incorporating BRPs can enhance assessment reliability and deliver robust parameters for fisheries management.

Although there are several studies evaluating catfish fishing resources in the Amazon basin (Petrere et al. 2004, Santanna et al. 2014, Doria et al. 2012) and the Pantanal (Mateus & Estupiñán 2002, Penha & Matheus 2004, Penha & Matheus 2007) and some species of Characiformes (Peixer et al. 2007), this study represents the first application of CMSY to assess Neotropical freshwater fish stocks. Previous applications focused predominantly on marine stocks, with limited use in European freshwater species (Fitzgerald et al. 2018, Andrašunas et al. 2022), Bangladesh (Khatun et al. 2023), and Indonesia (Nugroho & Uehara 2024). Given the unique dynamics of Amazonian ecosystems, future studies could complement this approach with length-based methods—such as LBI (Length-Based Indicators) (Froese 2004, Cope & Punt 2009) and LBSPR (Length-Based Spawning Potential Ratio) (Hordyk et al. 2015)—to validate the robustness of our findings.

## 5. CONCLUSIONS AND RECOMMENDATIONS

The results of this study, employing catch-based approach (CMSY), led to the following conclusions: a) Piracatinga was not overexploited during the period of highest fishing intensity (1996–2012) in the upper and middle Solimões River; b) The method suggest that Piracatinga fisheries can be considered sustainable, provided they are conducted without endangering dolphin, caiman, or other wildlife species by using them as bait; c) The CMSY method demonstrated sufficient accuracy for use in tropical fishery management, though we recommend cross-validation with length-based methods or regional comparisons to enhance robustness; d) Implementation of a catch-quota-based management model (e.g., *Quota Catch Transfer*, QCT) could mitigate potential growth overfishing scenarios, particularly when combined with complementary measures such as size regulations; e) Fisheries management strategies should also consider their migrations to ensure stock sustainability.

### Recommendations for Reopening the Piracatinga Fishery

For future management of Piracatinga fisheries, we advocate the adoption of catch-quota systems, specifically: Total Allowable Catch (TAC): A biomass-based percentage derived from MSY estimates to regulate fishing seasons and optimize regional fishery efficiency (Copes 1986). And Transferable Catch Quotas (TCQ): Allocation of TAC fractions to fishers, permitting usage rights or trade (Cavalcante & Furtado-Neto 2012). For Piracatinga, TCQ implementation would improve fleet economic efficiency, enable long-term catch projections, and incentivize fishers’ engagement in compliance and monitoring. Given the highly selective nature of Piracatinga fishing methods, bycatch of low-value or undersized species would be negligible.

## 6. AUTHOR CONTRIBUTIONS

Alfredo Perez Lozano: original draft writing, acquisition of data, review, and editing. Donald Charles Taphorn: analysis and interpretation of data, critical review and editing.

## 7. DATA AVAILABILITY STATEMENT

The entire data set that supports the results of this study was published in the article itself (Table I).

## 8. ACKNOWLEDGMENTS

The authors would like to extend their gratitude to the team at the SINCHI Institute in Colombia for all the support they received in carrying out this research. This work was carried out with the logistical and infrastructure resources of the Instituto de Ciências Exatas e Tecnologia of the Universidade Federal do Amazonas (UFAM/ICET), Brazil, and financial support from the Coordenação de Aperfeiçoamento de Pessoal de nível Superior (CAPES) in Brazil.

## REFERENCES

Agudelo E. 2007. La actividad pesquera en la zona suroriental de la Amazonía colombiana: una descripción de la captura y comercialización de los bagres transfronterizos. [Tesis de Doctorado]. Universidad Autónoma de Barcelona. Barcelona, España, 260 p. avalilable from https://ddd.uab.cat/record/142475

Agudelo E, Joven-León AV, Bonilla CA, Petrere Junior M, Peláez M, & Duponchelle F. 2013. Breeding, growth and exploitation of *Brachyplatystoma rousseauxii* Castelnau, 1855 in the Caquetá River, Colombia. Neotrop. Ichthyol.11:(3) 637–647.

Agudelo E, Salinas Y, Sánchez CL, Muñoz–Sosa DL, Alonso JC, Arteaga ME, Rodríguez OJ, Anzola NR, Acosta LE, Núñez M, & Valdés H. 2000. Bagres de la Amazonia colombiana: Un recurso sin fronteras. Fabré NN, Donato JC, & Alonso JC. (Eds.). Instituto Amazónico de Investigaciones Científicas SINCHI. Programa de Ecosistemas Acuáticos, Bogotá, Colombia 252 p.

Andrašunas V, Ivanauskas E, Švagždys A, & Razinkovasbaziukas A. 2022. Assessment of Four Major Fish Species Stocks in the Lithuanian and Russian Parts of Curonian Lagoon (Se Baltic Sea) Using Cmsy Method. Fishes 7: (1) 9. 10.3390/fishes7010009.

Barthem RB & Goulding M. 2007. Um ecossistema inesperado: A Amazônia revelada pela pesca. Amazon Conservation Association (ACA), Sociedade Civil Mamirauá. Tefé, Amazonas, Brasil, 241 p.

Berkson J, Barbieri L, Cadrin S, Cass-Calay SL, Crone P, Dorn M, Friess C, Kobayashi D, Miller TJ, Patrick WS, et al. 2011. Calculating Acceptable Biological Catch for Stocks That Have Reliable Catch Data Only (Only Reliable Catch Stocks). Noaa Technical Memorandum NMFS-SEFSC-616.

Bonilla CA, García A, Agudelo, E, Gómez-Hurtado G, Vargas G, & Duponchelle F. 2022. Life history trait variations and population dynamics of *Calophysus macropterus* (Siluriformes: Pimelodidae) in two river systems of the Colombian and Peruvian Amazon. Neotrop. Ichthyol. 20: (1), e210082. 10.1590/1982-0224-2021-0082.

Bonilla CA, Agudelo E, Acosta-Santos A, Gómez G, Ajiaco-Martínez RE, & Ramírez-Gil H. 2011. Calophysus macropterus. In: Lasso CA, Agudelo E, Jiménez-Segura LF, Ramírez-Gil H, Morales-Betancourt MA, & Ajiaco-Martínez R E (Eds). Catálogo de losRecursos Pesqueros Continentales de Colombia. Instituto de Investigación de Recursos Biológicos Alexander von Humboldt (IAvH). Bogotá, p. 432–435.

Bosch NE, Monk J, Goetze J, Wilson S, Babcock RC, Barrett N, Clough J, Currey-Randall LM, Fairclough DV, Fisher R, et al. 2021. Effects of human footprint and biophysical factors on the body-size structure of fished marine species. Biol. Conserv. 36: e-13807. 10.1111/cobi.13807

Brum SM, Da Silva VMF, Rossoni F, & Castello L. 2015. Use of dolphins and caimans as bait for Calophysus macropterus (Lichtenstein,1819) (Siluriformes: Pimelodidae) in the Amazon. J. Appl. Ichthyol. 31: 675–680. 10.1111/jai.12772

Carruthers TR, Punt AE, Walters CJ, Maccall A, Mcallister MK, Dick EJ, & Cope J. 2014. Evaluating methods for setting catch limits in data-limited fisheries. Fish. Res. 153: 48–68. 10.1016/j.fishres.2013.12.014

Cavalcante PPL & Furtado-Neto MAA. 2012. Implementation of individual transferable quotas and compulsory landing of live lobsters as a fishery management strategy. Arq. Cienc. Mar 45:(2) 49–59.

CCI. 2010. Pesca y Acuicultura de Colombia 2009. Corporación Colombiana Internacional. Bogotá

CCI. 2011. Pesca y Acuicultura de Colombia 2011. Corporación Colombiana Internacional. Bogotá.

Cope JM & Punt AE. 2009. Length-based reference points for data-limited situations: applications and restrictions. Mar. Coast. Fish. 1:(1) 169–186. 10.1577/C08-025.1

Copes P. 1986. A critical review of the individual quota as a device in fisheries management. Land Econ. 62:(3) 278–291.

De Oliveira-Dias J, Zanchi FB, Zacardi DM, Oliveira LS, De Vargas Schons S, & Sousa RGC. 2024. Population structure and reproductive indicators of the Surubim *Pseudoplatystoma punctifer* (Siluriformes, Pimelodidae) in the São Miguel River, Amazon basin, Brazil. J. Fish Biol. 104:(6) 1764 –1774. 10.1111/jfb.15714

Del Águila-Chávez J, Ríos-Pérez C, & Ríos-Ramírez R. 2023. Analysis of fishery landings of mota *Calophysus macropterus* in the Peruvian Amazon. Rev. BioCiencias 10: e-1542. 10.15741/revbio.10.e1542

Deza-Taboada SA, Bazán-Albites RS, & Culquichicon ZG. 2006. Bioecología y pesquería de *Pseudoplatystoma fasciatum* (Linnaeus, 1766; Pisces), doncella, en la región Ucayali. Folia Amazónica 14: 5–18. 10.24841/fa.v14i2.143

Doria CR, Ruffino ML, Hijazi NC, & Cruz RT 2012. A pesca comercial na bacia do rio Madeira no estado de Rondônia, Amazônia brasileira. Acta Amazônica 42: 29–40. 10.1590/S0044-59672012000100004

Drescher M, Perera AH, Johnson CJ, Buse LJ, Drew CA, & Burgman MA. 2013. Toward rigorous use of expert knowledge in ecological research. Ecosphere 4:(7) 1–26. 10.1890/ES12-00415.1

Fabré, N.N. and Alonso, J.C. 1998. Recursos Ícticos no alto Amazónas: Sua importância para as populações ribeirinhas. Bol. Mus. Paraense Emilio Goeldi, ser. Zool. 14: (1) 19–55.

FAO. 1995. Codigo de conducta para la pesca responsable. available from: https://openknowledge.fao.org/server/api/core/bitstreams/f10c6aea-f09e-4465-aa0a-37eb70ba5684/content,

Fitzgerald CJ, Delanty K, & Shephard S. 2018. Inland fish stock assessment: Applying data-poor methods from marine systems. Fish. Manage. Ecol. 25: (4) 240–252.10.1111/fme.12284

Franco D, Sobrane-Filho S, Martins A, Marmontel M, & Botero-Arias R. 2016. The piracatinga, *Calophysus macropterus*, production chain in the Middle Solimões River, Amazonas, Brazil. Fish. Manage. Ecol. 23:(2) 109–118. 10.1111/fme.12160

Freitas TMS & Montag LFA. 2019. Population and reproductive parameters of the red-tailed catfish, *Phractocephalus hemioliopterus* (Pimelodidae: Siluriformes), from the Xingu River, Brazil. Neotrop. Ichthyol. 17: e190015. 10.1590/1982-0224-20190015

Froese R. 2004. Keep it simple: Three indicators to deal with overfishing. Fish Fish. 5: 86–91. 10.1111/j.1467-2979.2004.00144.x

Froese R, Demire LN, Coro G, & Winker H. 2019. A Simple User Guide for CMSY+ and Bsm (CMSY_2019_9f.R). Oceanrep: Kiel, Germany.

Froese R, Demirel N, Coro G, Kleisner KM, & Winker H. 2017. Estimating fisheries reference points from catch and resilience. Fish 18: 506–526. 10.1111/faf.12190

Froese R, Garilao C, Winker H, Coro G, Demirel N, Tsikliras A, Dimarchopoulou D, Scarcella G, & Sampang–Reyes A. 2016. Exploitation and Status of European Stocks. Updated Version; Oceanrep: Kiel, Germany, 2016. available from: https://oceanrep.geomar.de/34476/

Froese R, Winker H, Coro G, Demirel N, Tsikliras AC, Dimarchopoulou D, Scarcella G, Palomares MLD, Dureuil M, & Pauly D. 2022. Estimating stock status from relative abundance and resilience. Ices J. Mar. Sci. 77:(2) 527–538. 10.1093/icesjms/fsz230

García A, Alonso J, Carvajal F, Moreau J, Nuñez J, Renno JF, Tello S, Montreuil V, & Duponchelle F. 2009. Life-history characteristics of the large Amazonian migratory catfish *Brachyplatystoma rousseauxii* in the Iquitos region, Peru. J. Fish Biol. 75: 2527–2551. 10.1111/j.1095-8649.2009.02444.x

Garcia A, Ruíz L, Vargas G, Sánchez H, Tello S, & Duponchelle F. 2017.Alimentación natural de la Mota *Calophysus macropterus* (Lichtenstein, 1819), En ambientes de la Amazonía Peruana. Folia Amazonica 26:(1) 29–36. 10.24841/fa.v26i1.416

Gómez C, Trujillo F, Diaz-Granados MC, & Alonso JC. 2008. Capturas dirigidas de delfines de río en la Amazonia para la pesca de la mota (*Calophysus macropterus*): una problemática regional de gran impacto. En: Trujillo F, Alonso JC, Diaz-Granados MC, & Gomez C. (Eds.). Fauna acuática amenazada en la Amazonia colombiana: análisis y propuestas para su conservación. Fundación Omacha, Corpoamazonía, SINCHI, Fundación Natura. Bogotá, Colombia. p. 39–57.

Gulland J A. 1969. Manual of methods for fish stock assessment. Part 1. Fish population analysis. Fao Fish. Tech. Paper, No. 4, Rev. 1.

Gushchin AV & Shavrina IA 2018. Current state of commercial fish stock in estuaries in the southern part of the Baltic Sea as a consequence of anthropogenic impact. Reg. Ecol. 52: 54–64. 10.30694/1026-5600-2018-2-43-53

Hordyk A, Ono K, Valencia S, Loneragan N, & Rince J. 2015. A novel length-based empirical estimation method of spawning potential ratio (SPR), and tests of its performance, for small-scale, data-poor fisheries. Ices J. Mar. Sci. 72:(1) 217–31. 10.1093/icesjms/fsu004

INCODER. 2006. Boletín estadístico 2001-2005. Instituto Colombiano de Desarrollo Rural –pesquero. Bogotá Colombia.

INPA, 1999. Boletín estadístico pesquero 1997-1998. Bogotá, Colombia.

INPA, 2001. Boletín estadístico pesquero 1999-2000. Bogotá Colombia.

Iriarte V & Marmontel M. 2013. Insights on the use of dolphins (boto, *Inia geoffrensis*, and tucuxi, *Sotalia fluviatilis*) for bait in the piracatinga (*Calophysus macropterus*) fishery in the western Brazilian Amazon. J. Cetacean Res. Manage. 13:(2) 163–173.

Johannes R E. 1998. The case for data-less marine resource management: examples from tropical nearshore fisheries. Trends Ecol. Evol. 13:(6) 243–246.

Khatun M H, Zahangir MM, Akhter B, Parvej MR, & Liu Q. 2023. Clupeids in the kaptai reservoir, a blessing or a curse: estimation of fisheries reference points. Heliyon 9: e-13818. 10.1016/j.heliyon.2023.e13818

Martell S & Froese R 2013. A simple method for estimating Msy from catch and resilience. Fish Fish. 14:(4) 504–514. 10.1111/j.1467-2979.2012.00485.x

Martin TG, Burgman MA, Fidler F, Kuhnert PM, Low-Choy S, Mcbride M, & Engersen K. 2012. Eliciting expert knowledge in conservation science. Conserv. Biol. 26: (1) 29–38. 10.1111/j.1523-1739.2011.01806.

Mateus LAF & Estupiñán GMB. 2002. Fish stock assessment of piraputanga *Brycon microlepis* in the Cuiabá River basin, Pantanal of Mato Grosso, Brazil. Braz. J. Biol. 62:(1) 165–170. 10.1590/S1519-69842002000100018

Ministério do Meio Ambiente. 2014. Instrução Normativa Interministerial MMA-Mpa No.6, de 17 de julho de 2014. Diário Oficial da União. Brasília. Available from:https://www.icmbio.gov.br/cma/images/stories/Legislacao/Instru%C3%A7%C3%B5es_normativas/IN_mma_n6_de_17_de_julho_2014_Piracatinga.pdf

Newman D, Berkson J, & Suatoni L. 2015. Current methods for setting catch limits for data-limited fish stocks in the United States. Fish. Res. 164: 86–93. 10.1016/j.fishres.2014.10.018

Nugroho S & Uehara T. 2024 Navigating Crisis: Insights into the Depletion and Recovery of Central Java’s Freshwater Eel (*Anguilla spp*.) Stocks. Sustainability 16: e-1578. 10.3390/su1604157

Nuñez-Avellaneda M, Agudelo E, Alonso J C, & Escobar M D. 2007. Ecosistemas acuáticos. En: Murcia UG. (Ed.). Balance anual sobre el estado de los ecosistemas y el ambiente de la Amazonia colombiana. SINCHI. Bogotá, p. 71–91.

Pauly D. 1997. Small-scale fisheries in the tropics: marginality, marginalization, and some implications for fisheries management. Global Trends: Fisheries Management 20: 40–49.

Peixer JA, Catella AC & Petrere Júnior M. 2007. Yield per recruit of the pacu *Piaractus mesopotamicus* (Holmberg, 1887) in the Pantanal of Mato Grosso do Sul, Brazil. Braz. J. Biol. 67:(3) 561–567. 10.1590/S1519-69842007000300023

Penha J M F, Mateus LAF, & Barbieri G. 2004. Age and Growth of the Porthole Shovelnose. Catfish (*Hemisorubim platyrhynchos*) in the Pantanal. Braz. J. Biol. 64:(4) 833–840. 10.1590/S1519-69842004000500013

Penha JMF & Mateus LAF. 2007. Sustainable harvest of two large predatory Catfish in the Cuiabá River basin, northern Pantanal, Brazil. Braz. J. Biol. 67:(1) 81–89. 10.1590/S1519-69842007000100011

Pereira DL & Hansen MJ. 2003. A Perspective on Challenges to Recreational Fisheries Management: Summary of the Symposium on Active Management of Recreational Fisheries. North Am. J. Fish. Manage. 23:1276–1282.

Pérez A & Fabré N 2002. Aspectos Reproductivos de la Piracatinga *Calophysus macropterus* Lichtenstein, 1819 (Pisces: Pimelodidae) en la Amazonia Central, Brasil. Bol. Cen. Invest. Biol. 36:(3) 217–374.

Pérez A & Fabré NN. 2009. Seasonal growth and life history of the catfish *Calophysus macropterus* (Lichtenstein, 1819) (Siluriformes: Pimelodidae) from the Amazon floodplain. J. Appl. Ichthyol. 25:(3) 343–349. 10.1111/j.1439-0426.2008.01104.x

Petrere M, Barthem RB, Cordoba EA, & Gomez BC. 2004. Review of the large catfish fisheries in the upper Amazon and the stock depletion of Piraiba (*Brachyplatystoma filamentosum* Lichtenstein). Rev. Fish Biol. Fish. 14: 403–414.

Pons M, Cope JM, & Kell LT. 2020. Comparing performance of catch–based and length–based stock assessment methods in data– limited fisheries. Can. J. Fish. Aquat. Sci. 77:1026–1037. 10.1139/cjfas-2019-0276

Prince J, Victor S, Kloulchad V, & Hordyk A. 2015. Length based Spr assessment of eleven Indo-Pacific coral reef fish populations in Palau. Fish. Res. 171: 42–58. 10.1016/j.fishres.2015.06.008.

Santanna IR, Doria CRC, & Freitas CE. 2014. Pre-impoundment stock assessment of two Pimelodidae species caught by small-scale fisheries in the Madeira River (Amazon Basin – Brazil). Fish. Manage. Ecol. 21: 322–329.10.1111/fme.12082

Schaefer MB. 1954. Some aspects of the dynamics of populations important to the management of the commercial marine fisheries. Inter-Am. Trop. Tuna Comm. Bull. 1:27–56. Available from: https://aquadocs.org/handle/1834/21257

SEPEC. 2017. Análisis de la estructura de tallas de captura de las principales especies ícticas explotadas por las pesquerías artesanales de Colombia. Bogotá, Colombia

Sharma R, Winker H, Levontin P, Kell L, Ovando D, Palomares MLD, Pinto C, & Ye Y. 2021. Assessing the Potential of Catch–Only Models to Inform on the State of Global Fisheries and the UN’s SDGs. Sustainability 13:(11) e-6101. 10.3390/su13116101

Sparre P & Venema SC. 1997. Introduction to Tropical Fish Stock Assessment, Part 1. Fao Fish. Tech. Pap. No 306/1 Rev. 2. Roma, Italy.

Stergiou KI. 2002. Overfishing, tropicalization of fish stocks, uncertainty and ecosystem management: Resharpening Ockham’s razor. Fish. Res. 55:(1–3) 1–09. 10.1016/S0165-7836(01)00279-X

Tomas ARG, Eleuterio CLT & Velasco G. 2019. Life History Parameters of *Genypterus brasiliensis* (Teleostei: Ophidiidae), an Endemic Fisheries Resource of the Southwestern Atlantic. Ann. Acad. Bras. Cienc. 91: e20170793. 10.1590/0001-3765201920170793

Van Damme PA, Navia-Morató J, Marmontel M, Trujillo F, Sainz L, Echeverría A, & Córdova CL. 2023. Fisheries of the scavenger species *Calophysus macropterus*: a case study in the Bolivian Amazon. Neotrop. Hydrobiol. Aquat. Conserv. 4: (1) 73–96. 10.55565/nhac.khgu2735

